# Eye-specific drive and binocular suppression in mouse visual cortex during critical period development

**DOI:** 10.64898/2026.09.10.750218

**Authors:** Katya Tsimring, Kyle R Jenks, Yuma Osako, Mriganka Sur

**Affiliations:** Department of Brain and Cognitive Sciences, Massachusetts Institute of Technology, Cambridge, MA, 02139, USA; The Picower Institute for Learning and Memory, Massachusetts Institute of Technology, Cambridge, MA, 02139, USA

**Author notes:** These authors contributed equally (Katya Tsimring, Kyle R Jenks).

## Abstract

A key function of cortical sensory circuits is to integrate information from multiple sources to build a unified representation of the external environment. The binocular region of the mouse primary visual cortex (bV1) is a valuable model for studying sensory integration, as visual response properties of inputs from the contralateral (contra) and ipsilateral (ipsi) eye onto bV1 neurons become matched to one another over development. While the alignment of visual responses from each eye during this developmental critical period has been well-characterized, it remains unclear how this interocular alignment influences, and is influenced by, the binocularly driven responses of bV1 neurons. Here, we recorded the monocular and binocular visual response properties of layer 2/3 bV1 neurons by performing chronic two-photon calcium imaging at multiple time points spanning the ocular-dominance critical period. We found that ipsi driven responses strengthened in bV1 neurons alongside a concurrent increase in binocular suppression. Binocular tuning properties were more strongly aligned with contra responses than with ipsi responses throughout development but became progressively better matched to ipsi responses. Additionally, the explained variance of ipsi input to predicting binocular tuning increased. In chronically tracked bV1 neurons, the preferred orientation of ipsi eye driven responses was less stable than that of contra eye driven responses, and in well-tuned stable binocular neurons, ipsi and contra preferences became progressively better matched. Using population-based decoding analyses, we found that while monocular and binocular visual encoding remained stable, there was increased generalizability between visual encoding by binocular and ipsi eye driven responses. Overall, our data suggest a reciprocal interaction between ipsi development and binocular responses, such that ipsi visual inputs dynamically structure excitatory as well as putative inhibitory drive onto binocular neurons during the critical period to generate the circuitry that implements integrated binocular vision.

## Introduction

Immature sensory circuits undergo experience-dependent reorganization during critical periods of elevated synaptic plasticity, which facilitates the creation of a coherent representation of the sensory environment. The primary visual cortex (V1) is a valuable model for studying the experience-dependent maturation of sensory circuits, as the integration and proper alignment of visual information from the two eyes onto binocular neurons in V1 requires visual experience (Wang et al., 2010, 2013; Espinosa and Stryker, 2012; Chang et al., 2020). Depriving one or both eyes of visual experience during this critical period impairs this alignment (Levine et al., 2017), and has been shown to have long lasting effects on binocular vision and depth perception (Prusky and Douglas, 2003; Williams et al., 2015; Scholl et al., 2017). While the developmental alignment of single-eye visual responses has been well-characterized (Jenks and Shepherd, 2020; Tan et al., 2020, 2022), it remains unclear how this relates to the development of visual response during normal, binocular viewing in single neurons or at a population level over the critical period.

Much of our knowledge on the experience-dependent maturation of binocular neurons in V1 comes from rodent studies due to the wide array of genetic tools, accessibility, and similarity of the primary visual pathway across mammals. At the onset of eye-opening at postnatal day 14 (p14), neurons in the binocular region of the rodent visual cortex (bV1) predominately respond to the contralateral (contra) eye (Smith and Trachtenberg, 2007; Jenks and Shepherd, 2020; Tan et al., 2021, 2022), and exhibit adult levels of selectivity for receptive fields, orientations, and directions (Rochefort et al., 2011; Hoy and Niell, 2015; Tan et al., 2022) – suggesting that the development of contra eye driven responses is experience-independent. While the fraction of bV1 neurons driven by the ipsilateral (ipsi) eye rapidly increases within a week of eye opening, the ipsi eye driven responses are poorly tuned in comparison to the contra eye (Smith and Trachtenberg, 2007; Tan et al., 2021, 2022). Additionally, bV1 neurons that respond to both the contra and ipsi eye at the start of the critical period (∼p21) have mismatched orientation preferences to each eye, indicating poor binocular integration in this early circuit (Wang et al., 2010, 2013; Tan et al., 2020, 2022). By the end of the critical period (∼p35), the ipsi and contra preferences in bV1 neurons are well-matched and there is a significant improvement in the tuning of ipsi eye driven responses (Wang et al., 2010, 2013; Tan et al., 2020, 2022). Together, these results suggest that the alignment of visual information from the contra and ipsi eye occurs through the developmental refinement of ipsi inputs onto bV1 neurons.

Despite the body of work done to characterize the visual properties of ipsi and contra eye responses in bV1 neurons, it remains unclear how these changes in monocular visual responses compare to visual responses driven by both eyes simultaneously during normal binocular viewing. While most findings hypothesize that initially weak ipsi responses align to contra responses through experience-dependent plasticity, a recent study in ferrets found that binocular visual properties in neurons are separate from and more stable than contra responses during the critical period (Chang et al., 2020). This study negates the view that developmental binocular responses simply reflect contra responses and instead suggests that eye-specific alignment should be studied in the context of binocular experience driving plasticity. This framework is intuitive given that an animal’s natural state during development is simultaneous, rather than separate, viewing through each eye and that binocular vision is known to be a nonlinear combination of monocular responses (Longordo et al., 2013; Zhao et al., 2013). Thus, it is crucial to study the simultaneous development of monocular and binocular responses to determine how visual inputs are truly integrated and refined over the critical period.

In this study, we performed two-photon calcium imaging of layer 2/3 neurons in mouse bV1 every 5 days from the start (∼p22-24) to the end (∼p32-34) of the critical period and characterized tuning properties during monocular and binocular presentations of visual stimuli. We found that at a population level and in chronically tracked neurons, ipsi and contra eye driven responses exhibited distinct developmental profiles that impacted binocular responses and tuning selectivity. Consistent with earlier work (Tan et al., 2020, 2022), ipsi responses strengthened over the critical period, while binocular responses surprisingly decreased relative to monocular viewing indicating binocular suppression. We further found that binocular tuning preferences were strongly correlated with contra eye driven responses at all timepoints, unlike in ferrets, and that contra eye driven preferences were comparatively stable to binocular driven preferences across development. Ipsi responses, on the other hand, progressively stabilized and became aligned to the contra tuning preferences in orientation selective bV1 neurons. Finally, using population-based analyses, we found that generalizability of decoding between the ipsi eye and binocular vision increased over development. Our findings thus suggest that while binocular visual responses are predominately driven by contra inputs throughout the critical period, ipsi visual encoding gradually improves coinciding with an increase in binocular suppression, putatively through increased ipsi input onto inhibitory neurons.

## Results

### Chronic imaging of monocular and binocular visual responses in layer 2/3 excitatory neurons of the mouse bV1

To measure monocular and binocular visual responses in neurons during the critical period for orientation matching in bV1, we performed *in vivo* two-photon calcium imaging of excitatory layer 2/3 bV1 neurons expressing GCaMP6s and mRuby2. We imaged approximately every 5 days for a 10 day period; p22-24 (D1), p27-29 (D5), and p32-34 (D10) (**Figure 1A**) (Wang et al., 2010; Tan et al., 2020). We placed a window over bV1 using stereotaxic coordinates (see **Methods**) and validated the location using intrinsic optical imaging (**Figure 1B**). During two-photon imaging, mice were head-fixed and presented with a 1 second high-contrast sinusoidal grating drifting in 1 of 8 directions (0-315°), interleaved with a 3 second gray screen. Each unique grating was shown at 3 different spatial frequencies 0.02, 0.04, and 0.08 cycles per degree, totaling 24 unique direction-spatial frequency pairings each repeated 10 times. For each field of view, we measured responses during monocular viewing sessions (occluding either the eye ipsilateral or contralateral to the imaged hemisphere) and during a binocular viewing session. Overall, we recorded 4,506 neurons from 8 mice on D1, 2,517 neurons from 8 mice on D5, and 1,528 neurons from 6 mice on D10.

**Figure 1:**
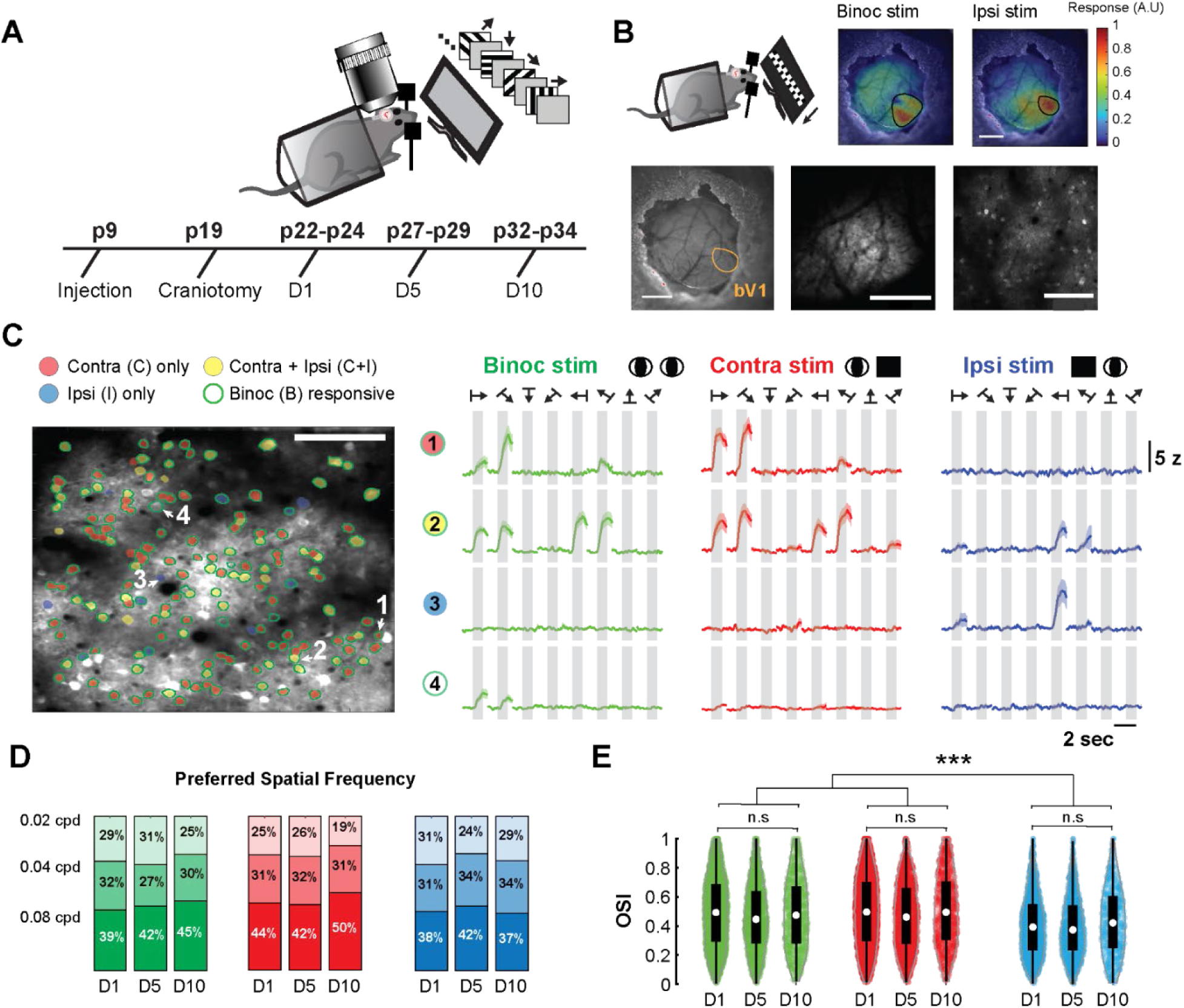
Chronic imaging of monocular and binocular visual responses in layer 2/3 excitatory neurons of the mouse bV1. **A.** Experimental timeline and recording setup**. B. (Top)** From left to right: example response amplitude map recorded during intrinsic optical imaging for binocular (binoc) stimulation and ipsi eye stimulation (scale bar 1mm). **(Bottom)** From left to right: Overlay of blood vasculature and area designated as the binocular region of V1 (scale bar 1mm). Brain surface in bV1 (scale bar 500 µm). Field of view imaged with two-photon (scale bar 100 µm). **C**. **(Left)** Average intensity projection of an example field of view recorded during two-photon calcium imaging on D1 (scale bar 100 µm). Neurons that were visually responsive are color coded based on their responsiveness to the contra eye, ipsi eye, and binoc stimulation. **(Right)** Trial-averaged calcium traces of 4 example neurons responding to each stimulation. Bolded lines denote the mean and shaded area denotes the standard error across the 10 trials. **D**. Proportion of neurons that preferred the 0.02, 0.04, and 0.08 cycles per degree (cpd) spatial frequency for each stimulation on D1, D5, and D10**. E.** Distribution of neuron’s OSI over days for the binocular, contra eye, and ipsi eye stimulation. While stimulation type had a significant effect on OSI (F(2, 12729), p=1.47e-37; 2-way ANOVA, main effect stimulation), the OSI did not significantly change from D1 to D10 across stimulation types (p = 0.8858, binoc stimulation; p=0.9838, contra stimulation; p=0.4983, ipsi stimulation; 2-way ANOVA and Tukey’s post hoc correction for stimulation and day).

We identified visual responses during binocular, contra eye, and ipsi eye viewing sessions by taking the mean response amplitude across the trials for each unique stimulus in each viewing session separately (**Figure 1C**). Overall, we found that 84.5% ± 3.7% of neurons were visually responsive in at least one viewing session on D1, 77.8% ± 4.9% on D5, and 68.7% ± 6.0% on D10. Our results are consistent with previous reports on the proportion of responsive neurons in bV1 during the critical period (Tan et al., 2020, 2022). To measure the tuning of bV1 neurons, we first identified a neuron’s preferred spatial frequency for the binocular, contra eye, and ipsi eye viewing session (**Figure 1D**). We then measured its orientation selectivity index (OSI) by taking the mean response amplitude across the 4 orientations at its preferred spatial frequency (**Figure 1E**). The binocular and contra OSI of neurons was significantly higher than ipsi at all timepoints, and mean OSI for both binocular and monocular viewing did not significantly change over development.

### Increase in ipsi drive coincides with increase in binocular suppression

To examine whether the bV1 population experienced a change in their eye-specific monocular responses across the critical period, we compared the proportion of neurons responding to the contra eye only (C), to the ipsi eye only (I), or to both the contra and ipsi eye (C+I neurons) (**Figure 2A**). In agreement with prior studies (Tan et al., 2020, 2022), we found that the proportion of ipsi responsive neurons significantly increased from D1 to D10 (D1: 10.9% ± 2.1% and D10: 24.0% ± 5.9%). The proportion of contra responsive neurons was consistent over development (D1: 48.7% ± 4.5% and D10: 49.9 ± 4.5%), while the proportion of C+I neurons decreased, but not significantly (D1: 40.9% ± 6.0% and D10: 26.0 ± 6.8%). To determine whether the relative drive from the contra and ipsi eye to C+I neurons changed from D1 to D10, we computed their ocular dominance index at each timepoint (ODI) and found that the initially contra dominated ODI progressively became more balanced over development (**Figure 2B**). Our results suggest that ipsi drive to bV1 strengthens over the critical period both through an increase in the number of neurons responsive only to the ipsi eye and increased ipsi drive to neurons responsive to both eyes.

**Figure 2:**
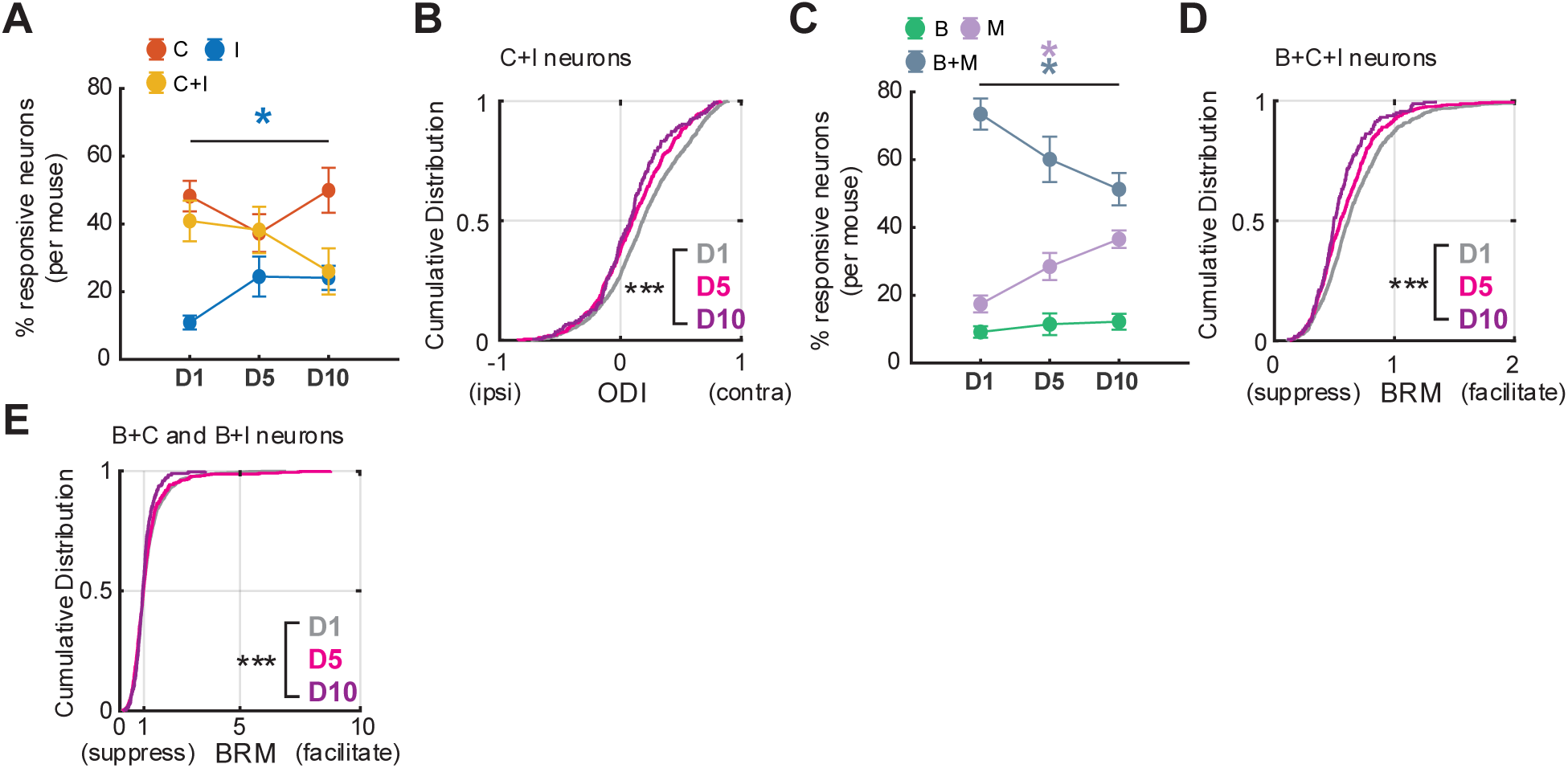
Increase in ipsi drive coincides with increase in binocular suppression. **A.** Proportion of neurons (mean per mouse) that responded only to the contra eye (C), only to the ipsi eye (I), or to both the contra and ipsi eye stimulation (C+I) by day. Proportion of I responsive neurons significantly changes from D1 to D10 (H(2)=7.0385, p=0.0296, % I responsive neurons; Kruskal-Wallis test). N = 8 mice (D1), 8 mice (D5), 6 mice (D10). **B.** Cumulative distribution of ODI in C+I responsive neurons. ODI shifts away from contra eye dominated responses from D1 to D10 (F(2,2017) = 19.63, p = 3.60*10^-9^; ANOVA). N = 1220 neurons on D1, 603 neurons on D5, 197 neurons on D10. **C.** Proportion of neurons that responded only to the binocular stimulation (B), only to the monocular stimulation (M), or to both the monocular and binocular stimulation (B+M) by day. Proportion of B+M neurons decreases while the proportion of M neurons increases (H(2)= 6.8755, p= 0.0321, % B+M responsive neurons; H(2)= 8.6591, p=0.0132, % M responsive neurons; Kruskal-Wallis test). **D.** Cumulative distribution of binocular response modulation (BRM) in B+C+I responsive neurons. Responses to binocular stimulation become suppressed compared to the sum of monocular stimulation from D1 to D10 (F(2,1742) = 15.9, p = 1.43*10^-7^, ANOVA). N = 1093 neurons on D1, 492 neurons on D5, 160 neurons on D10. **E.** Cumulative distribution of BRM in B+C and B+I responsive neurons also shows a decrease from D1 to D10 (F(2,2444) = 5.15, p = 5.8*10^-3^, ANOVA). N = 4506 neurons on D1, 2517 neurons on D5, 1528 neurons on D10. * p < 0.01, *** p < 0.0001.

How do these monocular changes relate to the fraction of responsive neurons during binocular viewing (**Figure 2C**)? The percentage of neurons that responded only to the monocular viewing sessions significantly increased from D1 to D10 (D1: 17.4% ± 2.5% and D10: 36.5% ± 2.6%), which coincided with a decrease in the percentage of neurons that responded to both the monocular and binocular viewing sessions on D10 (D1: 73.4% ± 4.6% and D10: 51.3% ± 4.8%). Single unit recordings in adult mice have shown that C+I neurons exhibit sublinear responses to binocular viewing compared to the sum of the monocular viewing sessions (Zhao et al., 2013). If this interocular suppression emerges during the critical period, it may explain the apparent loss of binocular responses for monocularly responding neurons. Indeed, in neurons that were responsive across all three viewing sessions (B+C+I), the peak response amplitude during the binocular session was, on average, lower than the linear sum of the contra and ipsi responses on D1 (**Figure 2D**) and decreased further across development. This increase in binocular suppression was also apparent in neurons that, in monocular sessions, responded to only the contra (B+C) or ipsi (B+I) eye (**Figure 2E**). Together, these findings demonstrate that binocular responses are increasingly suppressed over development, coinciding with an increase in ipsi drive to bV1 (Longordo et al., 2013; Zhao et al., 2013).

### Ipsi responses become matched to contra and binocular responses in orientation selective bV1 neurons

Previous studies have shown that bV1 neurons’ contra and ipsi eye inputs become aligned to the same orientation over the critical period (Wang et al., 2010, 2013; Chang et al., 2020; Tan et al., 2020, 2022). Whether this monocular alignment in mice is mirrored by alignment to an independent binocular representation (as shown in the ferret visual cortex; Chang et al., 2020) remains unclear. To address this question, we compared the contra eye, ipsi eye, and binocular tuning curves in neurons that were responsive to all three viewing sessions (B+C+I) (**Figure 3A**). It was recently shown that the conversion of C or I only responsive neurons to C+I responsive neurons is predicted by higher initial monocular OSI (Tan et al., 2020). Thus, we split responsive neurons by their OSI into well-tuned (OSI > 0.5) and poorly tuned (OSI < 0.5) neurons across each viewing sessions.

**Figure 3:**
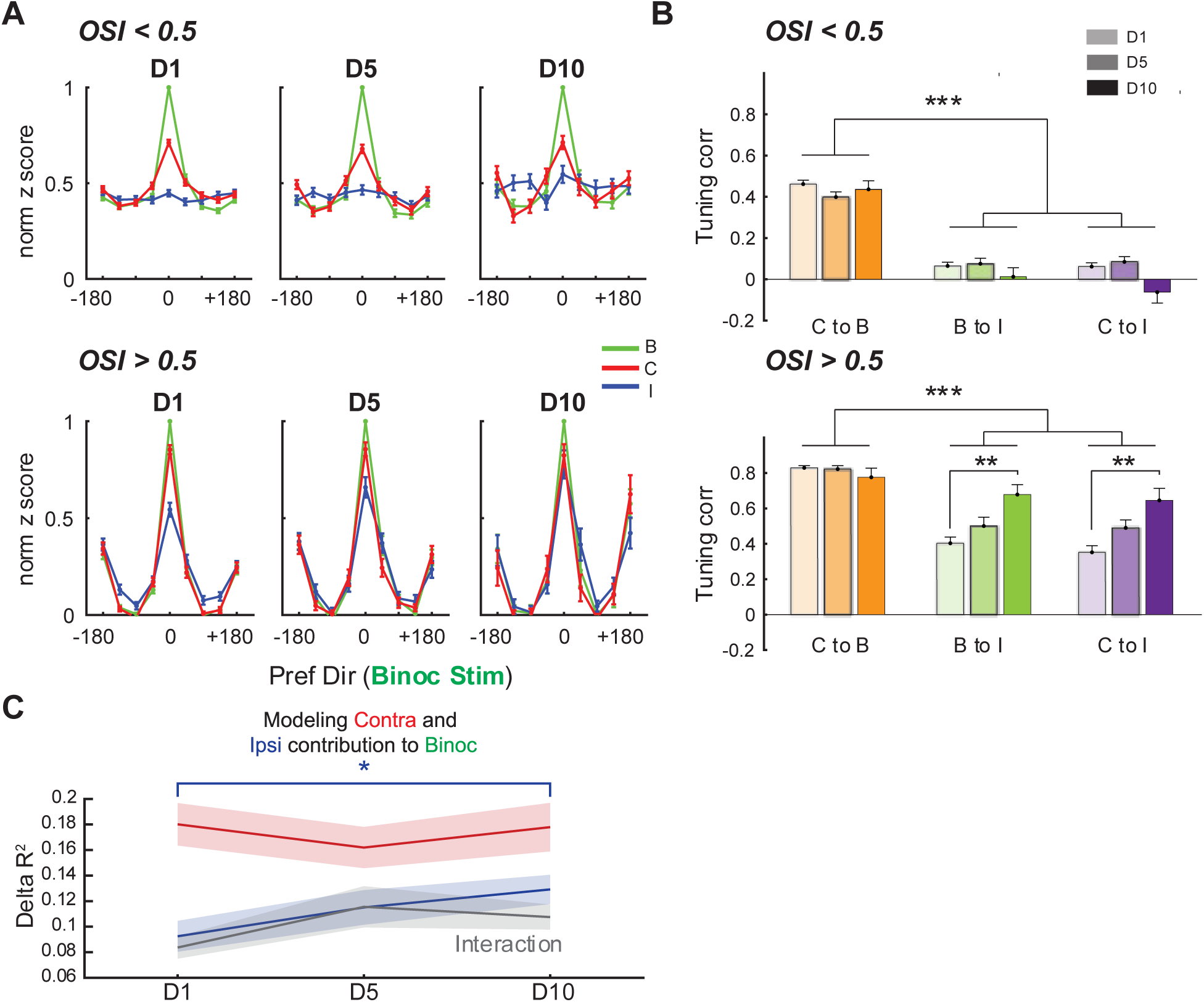
Ipsi responses become matched to contra and binocular responses in orientation selective bV1 neurons. **A.** Mean response amplitudes aligned to the binocular (binoc) stimulation’s preferred direction in neurons that were visually responsive across the three stimulations (mean ± s.e.m across neurons). Responses are normalized by the peak binocular response. (**Top**) neurons whose OSI for each stimulation is below 0.5. N = 395 neurons (D1), 204 neurons (D5), and 65 neurons (D10). (**Bottom**) neurons whose OSI for each stimulation is above 0.5. N = 159 neurons (D1), 66 neurons (D5), 25 neurons (D10). **B.** Correlation between the neuron’s tuning curves in each of the stimulations over days. (**Top**) Neurons whose OSI is below 0.5 for each stimulation session. Correlations between binocular (B) to contralateral eye (C), B to ipsilateral eye (I), and C to I tuning curves are significantly different but there is no effect of day on tuning curve correlations (F(2, 2386)=157.0838,p=8.16e-65, main effect stimulation comparison; F(2, 2386)= 2.8955, p**=**0.0555, main effect day; 2-way ANOVA). N = 641 neurons (D1), 317 neurons (D5), 109 neurons (D10). **(Bottom**) Neurons whose OSI is above 0.5 for each session. There is a significant interaction between stimulation comparison and day (F(2, 1169)= 5.4019, 2.60e-04; 2-way ANOVA, stimulation comparison and day interaction). B to I and C to I tuning correlation significantly increases from D1 to D10 (p=0.0067, B to I from D1 to D10; p= 0.0041, B to I from D1 to D10; 2-way ANOVA and Tukey’s post hoc correction for day and session pair). N = 452 neurons (D1), 175 neurons (D5), 51 neurons (D10). **C.** Linear regression models were used to predict the binocular tuning curve from the contra and ipsi tuning curves, as well as their interaction, for fields of view (FOVs) imaged across all 3 timepoints. The explained variance from each predictor is plotted per timepoint. Explained variance from contra did not change from D1 to D10 (F(1, 26) = 0.0090, p = 0.9253, linear mixed-effects model), while explained variance from ipsi increased significantly (F(1, 26) = 11.4289, p = 0.0023, linear mixed-effects model). There was a trend for increased explained variance from the contra:ipsi interaction (F(1, 26) = 3.5645, p = 0.0702, linear mixed-effects model). N = 14 FOVs (D1-D10). ** p < 0.01, *** p < 0.001.

In both well-tuned and poorly tuned neurons, the tuning curve during the binocular viewing session was more correlated (tuning/signal correlation; Cohen and Kohn, 2011) to the contra eye’s than to the ipsi eye’s tuning curve at each timepoint (**Figure 3B**). In the population of well-tuned neurons, but not of poorly tuned neurons, the ipsi eye tuning curve became better aligned to both the contra and binocular tuning curves over development. However, the ipsi tuning curve is not significantly more correlated to the binocular tuning curve than the contra tuning curve, or vice versa. Thus, the alignment of ipsi and contra tuning properties during the critical period occurs specifically for well-tuned bV1 neurons, consistent with previous findings (Tan et al., 2020), and ipsi alignment also occurs to the binocularly driven responses. The contra to binocular correlation, on the other hand, is already high at the start of the critical period and does not increase further. This suggests that binocular tuning in mice largely reflects contra driven responses across development.

The increase in ipsi input to bV1 coinciding with increased correlation of ipsi and binocular tuning led us to hypothesize that ipsi tuning would be increasingly predictive of binocular tuning over development. Therefore, we trained a linear regression model to predict the binocular tuning curve based on the contra and ipsi tuning curves, and separably measured the explained variance of contra, ipsi, and their interaction (contra:ipsi) across development (**Figure 3C**). We found that the ipsi-explained variance increased significantly across development, with a trend towards higher explained variance from the contra:ipsi interaction. The contra-explained variance remained high across development. These results show that ipsi tuning becomes more aligned with and predictive of binocular tuning over development.

### Ipsi input becomes more stable and aligned to contra eye preference over development

It was previously assumed that population level increased alignment of C+I orientation preference over development reflected the gradual alignment of neurons that were already C+I responsive at the start of the critical period. However, chronic imaging and tracking of single neurons has revealed that 60% of the initial, poorly aligned, C+I population at the start of the critical period are replaced by new, well aligned, C+I responsive neurons by critical period’s end (Tan et al., 2020). Whether the binocular responses are as labile as the monocular responses, and how monocular and binocular changes are related, is unclear. To address this, we selected neurons in our dataset that could be reliably identified on each of the three imaging days (**Figure 4A**). Where appropriate in our analysis, we also included neurons that could be identified in any two sequential timepoints.

**Figure 4:**
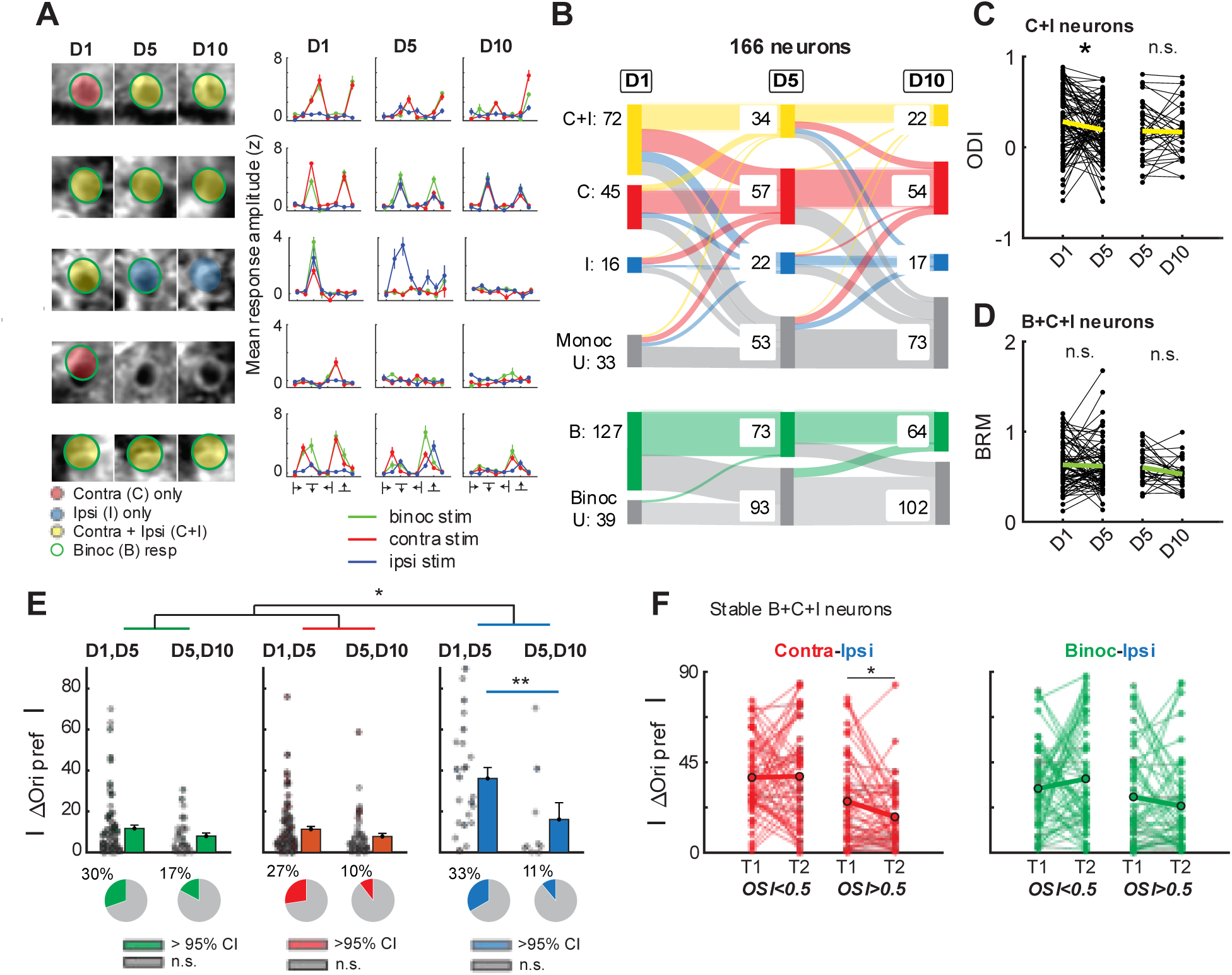
Ipsi input becomes more stable and aligned to contra eye preference over development. **A. (Left)** Five example neurons that were tracked from D1 to D10. Neurons are labeled by their response to the binocular (binoc), contra eye and ipsi eye viewing session**. (Right)** Trial-averaged mean response amplitudes for the binocular, contra eye, and ipsi eye viewing session from the sampled neurons. **B.** Flow chart tracking the fate of the **(Top)** eye-specific and **(Bottom)** binocular responses of 166 neurons that were tracked from D1 to D10**. C.** ODI in tracked neurons that were responsive to the contra and ipsi viewing session. ODI decreases in neurons from D1 to D5 (z = −2.3095, p = 0.0209, D1 to D5; z = −0.4531, p = 0.6505, D5 to D10; signed rank test). N = 89 neurons (D1-D5), 34 neurons (D5-D10). **D.** BRM in tracked neurons that were visually responsive across the three sessions. BRM does not change in tracked neurons (z = −0.3311, p = 0.7406, D1 to D5; z = −1.7298, p = 0.0837, D5 to D10; signed rank test) N = 72 neurons (D1-D5) and 27 neurons (D5-D10). **E. (Top)** Change in orientation preference from D1 to D5 and D5 to D10 in tracked neurons for the binocular, contra eye, and ipsi viewing session (mean ± s.e.m across neurons) for neurons with an OSI > 0.5. Neurons that experience significant changes in their orientation preference are color coded in green, red or blue for the respective session. **(Bottom)** Portion of neurons that exhibit significant changes above the 95% confidence interval. There is a significant interaction between stimulation session and day on orientation offset for tracked neurons (F(2,278)=3.6544, p=0.0271, 2-way ANOVA, stimulation session and day interaction). The offset in the preferred orientation for the ipsi viewing session decreases from D1-D5 to D5-D10 (p=0.0062, orientation offset for ipsi D1,D5 and D5,D10, 2-way ANOVA and Tukey’s post hoc correction for day and session). D1-D5: N = 79 neurons (binocular viewing), 95 neurons (contra eye viewing), 24 neurons (ipsi eye viewing). D5-D10: N = 29 neurons (binocular viewing), 48 neurons (contra eye viewing), and 9 neurons (ipsi eye viewing). **F.** Orientation offset between **(Left)** contra and ipsi or **(Right)** binoc and ipsi orientation preference for well-tuned and poorly tuned neurons that were B+C+I responsive at two sequential timepoints (T1, T2). For well-tuned B+C+I neurons, contra to ipsi offset decreased from T1 to T2 (z = 2.34, p = 0.019, signed rank test) N = 49 neurons (contra-ipsi, poorly tuned), 50 neurons (contra-ipsi, well-tuned), 53 neurons (binoc-ipsi, poorly tuned), 46 neurons (binoc-ipsi, well-tuned). * p < 0.05, ** p < 0.01, *** p < 0.001.

We first assessed stability in the monocular and binocular responsiveness of tracked neurons. Monocular responses of bV1 neurons were highly unstable from D1-D10 (**Figure 4B**), with only 38% of responsive neurons (50/133) maintaining their eye-specific response. Binocular responses were also unstable, with only 56% (70/127) of neurons maintaining their binocular response from D1-D10. In neurons that remained responsive to contra and ipsi viewing sessions (C+I) from D1 to D5, the ocular dominance shifted away from the contra eye (**Figure 4C**), suggesting that while turnover may account for most of the increase in ipsi responsiveness over development, there is also increased ipsi input to a subset of stable C+I responsive neurons. We did not observe a significant change in the BRM between timepoints for stable B+C+I neurons (**Figure 4D**).

For neurons that maintained binocular and/or contra responsiveness from D1 to D5, orientation preference remained relatively stable (**Figure 4E**: binocular: 11.7^°^ ± 1.6^°^, contra eye: 11.3^°^ ± 1.3^°^). Shifts in ipsi eye preference, on the other hand, were significantly larger than those for the contra eye and for binocular preference from D1-D5 (ipsi eye: 36.1^°^ ± 5.3^°^). From D5 to D10, however, the shifts in the ipsi eye preference were not significantly different than the shifts for contra and binocular viewing suggesting ipsi input had become more stable. To determine how shifts in monocular and binocular orientation preferences impacted orientation matching of ipsi input to the other viewing sessions, we examined how the offset in orientation preference for stable B+C+I neurons changed between two sequential timepoints (D1 to D5 or D5 to D10) (**Figure 4F**). We hypothesized that if binocular viewing drove ipsi alignment, then binocular to ipsi offset would decrease for stable B+C+I neurons. However, we instead found that ipsi input became more aligned to the contra orientation preference, and that this alignment was specific to well-tuned B+C+I neurons. Our results suggest that unlike in more binocular species such as ferret, binocular vision does not provide a separate, more stable, template for monocular responses to align to.

### Network level reorganization underlies developmental shift in visual encoding

We next addressed how the developmental changes we observed at a single neuron level impacted the representation of visual information at a network level. We used a 5-fold cross validated support vector machine (SVM) decoder to predict the stimulus orientation (**Figure 5A**) presented at a given trial using the mean response amplitudes of randomly sampled subsets of neurons. We found that the decoding accuracy for all viewing sessions remained stable for predicting orientation throughout development (**Figure 5B**). We hypothesized that developmental alignment of ipsi tuning to contra and binocular orientation preferences ensures that a given stimulus is encoded similarly regardless of which eye it is viewed through. Thus, we hypothesized that decoding between viewing sessions would become increasingly generalizable across development (**Figure 5A**), such that a decoder trained only on binocular or contra viewing would have improved decoding for ipsi viewing on D10 compared to D1. Indeed, we found that for a decoder trained on data from binocular viewing and tested on data from ipsi viewing the generalizability of the decoding increased significantly from D1 to D10 (**Figure 5C**). While there was a similar trend for decoding trained on contra data and tested on ipsi data, the result did not reach significance. Thus, over the course of the critical period we find that ipsi eye responses become more aligned to binocular responses at the network and single neuron level.

**Figure 5:**
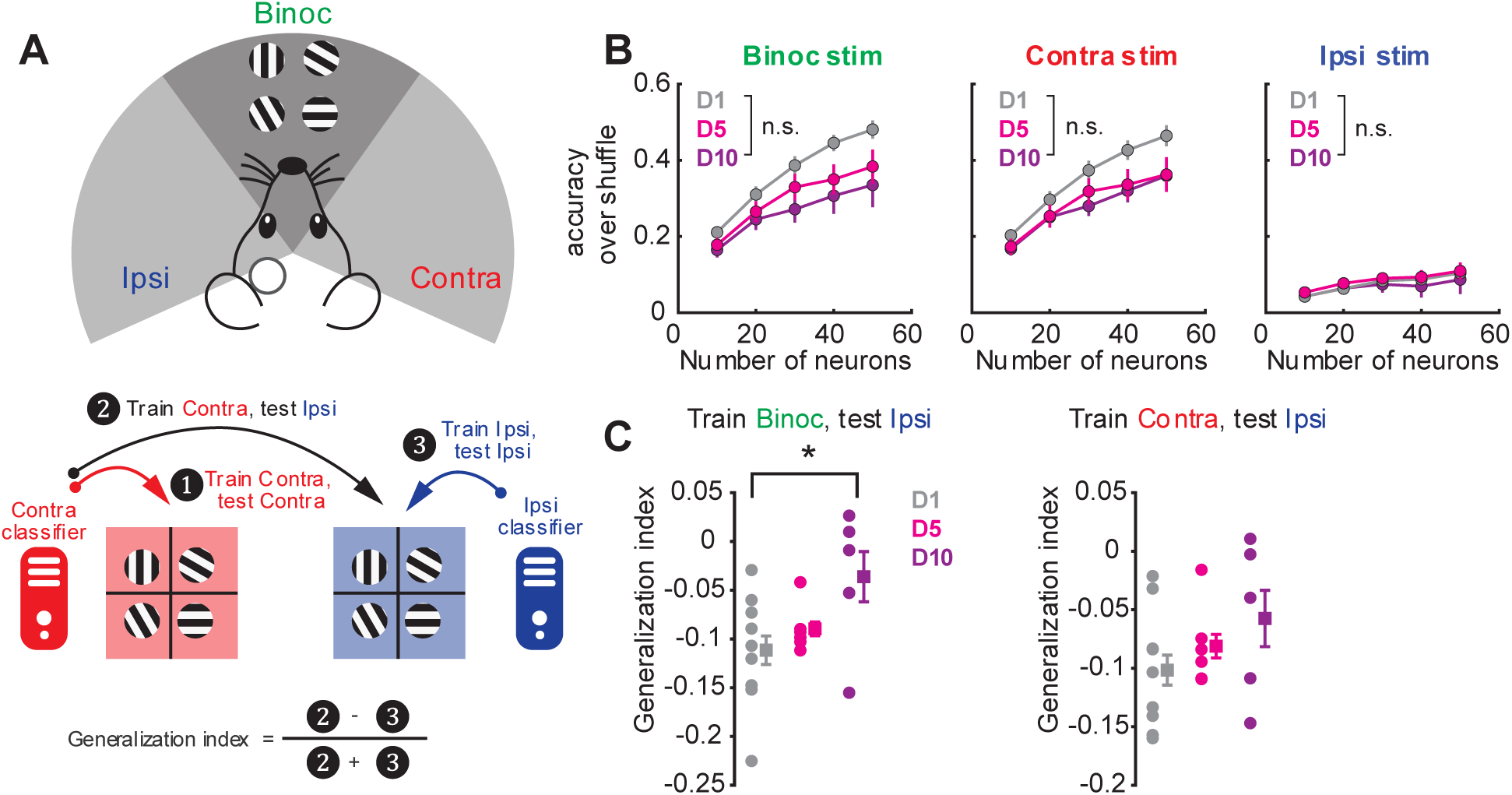
Network level reorganization underlies developmental shift in visual encoding. **A.** Schematic of SVM decoder used for predicting orientation on a given trial. SVM decoders were trained with different population sizes (10-60 neurons) randomly sampled from each field of view in each viewing session (Binocular (binoc), contra, ipsi). Decoders were then tested within the same viewing session they were trained or tested against another viewing session (but the same population of neurons) in order to compute the generalization index. **B.** Decoding compared to shuffle for binoc, contra, and ipsi viewing sessions on D1, D5, and D10. There was no improvement within viewing sessions between D1, D5, and D10. **C.** Generalization index quantifying decoding accuracy between viewing sessions. Decoders trained on binocular viewing and decoding stimulus identity during ipsi viewing had significantly increased generalizability from D1 to D10 (p = 0.0252) with a significant positive going slope from D1 to D10 showing improvement over the course of the critical period (r = 0.4981, p = 0.0246). Decoders trained on contra viewing and decoding stimulus identity during ipsi viewing did not have a significant increase in the generalization index (p = 0.1448) or positive going slope (r = 0.3496, p = 0.1302). For all neurons: N = 32 FOVs (D1), 23 FOVs (D5), and 18 FOVs (D10). For binocular responsive neurons: N = 27 FOVs (D1), 20 FOVs (D5), 11 FOVs (D10). For contra responsive neurons: 27 FOVs (D1), 20 FOVs (D5), 13 FOVs (D10). For ipsi responsive neurons: 26 FOVs (D1), 18 FOVs (D5), 8 FOVs (D10). * p < 0.05.

## Discussion

During the critical period for binocular vision, neurons in the mouse bV1 undergo developmental changes in their monocularly driven visual responses (Wang et al., 2010, 2013; Jenks and Shepherd, 2020; Tan et al., 2020). How these eye-specific changes are reflected in the binocular representation of visual information remains unclear. Our goal was to understand the relationship between visual responses generated by one or both eyes in bV1 neurons over development by performing chronic two-photon calcium imaging in awake mice during the critical period. By characterizing visual properties at the population level and in individual neurons, we demonstrate that ipsi eye strengthening coincides with increased suppression of binocular responses, and that the bV1 neural network undergoes reorganization to facilitate the generalizable encoding of visual information from the ipsi eye.

### Development of binocular sublinear integration

In agreement with prior studies that have recorded ipsi and contra eye driven responses of bV1 neurons (Tan et al., 2020, 2022), we found that there was an overall increase in ipsi drive to bV1 and a gradual alignment in the tuning preferences between the contra and ipsi eye in C+I neurons, specifically in orientation selective neurons. However, there was also a reduction in responses to binocular stimulation compared to monocular responses. Sublinear integration of binocular cortical responses is a phenomenon that is shared across species, in mice (Longordo et al., 2013; Zhao et al., 2013), in cats (Sengpiel and Vorobyov, 2005), in macaques (Dougherty et al., 2019; Mitchell et al., 2022), and in humans (Moradi and Heeger, 2009), and it is important for preserving binocular tuning and generating disparity in mouse bV1 (Longordo et al., 2013; Zhao et al., 2013). Our results thus demonstrate an important, unsuspected element of binocular convergence during development, whereby suppressive interactions co-evolve with ipsi strengthening.

How might binocular sublinear integration arise during the critical period? Previous studies have shown that cortical inhibition increases after eye-opening (Hensch et al., 1998; Feldman, 2000; Iwai et al., 2003; Lazarus et al., 2011). Since the increase in cortical inhibition coincides with the addition and strengthening of ipsi inputs (Smith and Trachtenberg, 2007; Faguet et al., 2009; Tan et al., 2021), it is possible that the inputs onto inhibitory neurons from the ipsi eye are also increasing and thus enhance ipsi eye driven feedforward inhibition onto binocular neurons. Indeed, interocular suppression through cortical inhibition has previously been reported in cats to reduce responses at the preferred orientation (Sengpiel and Vorobyov, 2005). Binocular suppression could also emerge prior to bV1 in the thalamus, as binocular responses in the thalamus have also been shown to integrate inputs sublinearly (Howarth et al., 2014; Huh et al., 2020); however, sublinear integration in bV1 neurons has been reported to be the result of inhibitory, not excitatory, drive (Longordo et al., 2013; Zhao et al., 2013). Future work will need to be done to examine the underlying source of binocular suppression over development, and whether it is causally related to the strengthening of thalamic and/or cortical ipsi inputs onto inhibitory neurons.

### Contra inputs set the tuning and alignment of binocular responses in mouse bV1

One of the prominent features we observed across development was the strong congruence between neural responses to the contra and binocular viewing sessions. We found that contra and binocular responses were aligned to similar orientations and highly correlated across the critical period. Furthermore, in chronically tracked neurons, the contra and binocular orientation preferences were stable at the beginning of the critical period and did not undergo further stabilization, unlike ipsi orientation preferences. Based on our findings, we propose that contra inputs provide a crucial backbone for driving the gradual integration, stabilization and alignment of ipsi inputs onto bV1 neurons during the critical period (Smith and Trachtenberg, 2007; Faguet et al., 2009).

Though our results are not surprising given the dominance of contra inputs in mouse bV1 (Gordon and Stryker, 1996; Cang et al., 2023), our findings differ from earlier work described in the ferret visual cortex, in which the binocular preference was *independent* of the contra and ipsi responses and served as the template for ipsi-contra eye alignment (Chang et al., 2020). The distinction in our findings can be attributed to species-related differences in the organization of the ferret and mouse visual cortex. The ferret V1 contains large-scale cortical columns for ocular dominance and orientation preference (Law et al., 1988; Chapman et al., 1996), whereas mouse V1 circuitry is more intermixed and localized in comparison (Espinosa and Stryker, 2012). Thus, it is possible that the modular organization of the ferret V1 facilitates an independent representation of binocular inputs via the amplified recurrence emerging from its lateral inputs. On the other hand, previous studies have shown that thalamic inputs on L4 neurons in bV1 exhibit binocular matching preceding that of intracortical inputs (Gu and Cang, 2016), suggesting a potential feed-forward model for ipsi-contra alignment in mouse V1.

### Impact of circuit level reorganization on visual encoding

The putative purpose of experience-dependent developmental plasticity is to modify synaptic connections to enhance sensory integration and encoding. We therefore hypothesized that the developmental dynamics in responses we observed at the single cell level altered the encoding of visual information at the population level. While decoding of the stimulus from neuronal responses did not improve over development for monocular or binocular responses, we found increased generalizability between the binocular and ipsi viewing sessions suggesting improved integration between monocular and binocular encoding and supporting our findings at the single neuron level that ipsi tuning becomes better aligned to and predictive of binocular tuning. While we did not see a significant change in the decoding of binocular responses, it is important to note that grating stimuli may be too simple to distinguish a developmental change in visual encoding of binocular representations, as the parameter space we explored does not cover the rich statistics of naturalistic stimuli (Kayser et al., 2004; Rikhye and Sur, 2015). Thus, it will be critical in future work to record neuronal responses to natural images to study how population level encoding changes over the critical period.

## Author Contributions

This project was conceptualized by M.S., K.T., and K.J. All data collection was performed by K.T. Analysis was performed by K.T., K.J., and Y.O. Writing was done by K.T. and K.J. with input from M.S. and Y.O. All authors read and approve the final manuscript.

## Acknowledgements

This work was supported by NIH grants R01MH126351 (M.S.), R01EY028219 (M.S.), F31EY033649 (K.T.), and F32EY032756 (K.J.). In addition, this work was supported by the Picower Institute Innovation Fund (M.S.), Japan Society for the Promotion of Science (JSPS) Overseas Research Fellowships (Y.O.), and The Uehara Memorial Foundation Postdoctoral Fellowship (Y.O.). The authors thank Taylor Johns and other members of the Sur lab for their help and support.

## Competing interests

The authors declare no competing interests.

## Methods

### Experimental Model and Subjects

All procedures performed in this study were approved by the Massachusetts Institute of Technology’s Animal Care and Use Committee and conformed to the Guide for the Care and Use of Laboratory Animals published by the National Institutes of Health. Male and female wild-type C57BL/6j mice were used in study. Mice were group-housed (no more than five mice per cage) with a standard light/dark cycle of 12/12 hours with access to food and water ad libitum.

### Stereotaxic Surgery Procedures

#### Viral injection

Postnatal (p) day 9-10 pups were anesthetized with isoflurane (3% for induction, 1-1.5% for maintenance) while maintaining a body temperature of 37.5 degree C° using a heating pad (ATC2000, World Precision Instruments) and placed on a stereotaxic apparatus (Kopf). Pre-operative slow-release buprenorphine (1mg/kg, intraperitoneal injection) and post-operative meloxicam (5mg/kg, subcutaneous injection) were provided to mice before surgery. Once the appropriate level of anesthesia was achieved, fur was removed from the surgical site with Nair. Skin was cleaned with saline, betadine, and 70% ethanol three times with a cotton applicator. The scalp was then scored with a scalpel and the skin was folded back to expose the skull. Using stereotaxic coordinates for the left hemisphere of the binocular region of the visual cortex (0.5-1 mm anterior from the lambda suture and 3 mm lateral from the midline suture), 150 nL of a mixture of AAV9.CaMKII.Cre (105558-AAV9, Addgene) diluted 1:4 in AAV1-hSyn1-Flex-mRuby2-GSG-P2A-GCaMP6s-WPRE-SV40 (68720-AAV1, Addgene) (Rose et al., 2016) was injected through the skull (QSI 53311, Stoelting) at 50 nL/min (0.2-0.25 mm below the dura to target layer 2/3) using a glass pipette with a 50 μm diameter tip. We repeated this procedure 1-2 times at injection sites that were 500 μm apart. Following the injections, the skin was sutured with internal stitches (Prolene 7.0), and the pups were returned to the dam cage to recover.

#### Craniotomy and Head-plate Implant

9-10 days after the viral injection, mice were anesthetized and prepared for surgery using the same procedure as in *Viral Injection.* Following retraction of the scalp, we drilled and carefully removed a 3 mm circular piece of skull over the binocular region of the primary visual cortex, centered at 1.5 mm anterior from the lambda suture line and 3 mm lateral from the midline suture, in the left hemisphere using a dental drill. Two 3-mm coverslips centered on a 5-mm coverslip (CS-3R and CS-5R, Warner Instruments) were glued together with optical adhesive (NOA 61, Norland) and positioned over the craniotomy. The coverslip was then attached to the skull using Metabond (C&B Metabond, Parkell) mixed with a black ink pigment (Black Iron Oxide 18727, Schmincke). A custom designed stainless-steel head-plate was then positioned over the coverslip and attached to the skull with Metabond.

### Two-photon Imaging

Mice were head-fixed on a custom-built behavior rig and placed in a polypropylene tube to constrain movement. Two-photon imaging was done through the cranial window over the binocular region of the primary visual cortex in mice using resonant-galvo scanning with a Prairie Ultima IV two-photon microscopy system (Bruker). For all recordings, we used a 16x 0.8 NA objective (CFI LWD Plan Fluorite, Nikon) and an excitation wavelength of 920 nm using a Ti:Sapphire tunable laser (InSight X3+, Spectra-Physics). Imaging was done at a digital zoom of 2× with a 512×512 resolution and frame rate of ∼7.75Hz following 4 frame averaging. To target recordings to layer 2/3, we imaged fields of view (FOVs) at 150-200 μm below the surface. For each FOV, we had three separate viewing sessions: binocular viewing, contralateral eye viewing, and ipsilateral eye viewing. Mice were imaged every 4-5 days for a period of ten days, starting from p22-p24 to encompass the critical period for binocular orientation matching (Wang et al., 2010, 2013; Tan et al., 2020).

### Visual Stimulus for Two-Photon Imaging

An LCD monitor (11.6-inch, 60 Hz refresh rate, LONCEVON) was placed about 9 cm in front of the mouse to display the visual stimulus. The monitor covered ∼100 degrees in azimuth and ∼60 degrees in elevation and had a mean luminance of 35 cd/m^2^. During two-photon imaging, we presented awake mice with high-contrast sinusoidal gratings that drifted at 8 different directions separated by 45-degree intervals with a temporal frequency of 2Hz, and spatial frequencies of 0.02, 0.04, and 0.08 cycles per degree (Psychophysics Toolbox in MATLAB). Ten trials of each direction/cycles per degree combination were presented in a pseudorandom order. Each trial was followed by a 3 second gray screen period (inter-stimulus interval).

### Optical Imaging

To determine whether the injection site was within the binocular region of the primary visual cortex, optical imaging was performed on 5 out of the 8 mice. Mice were head-fixed on a behavioral rig and lightly anesthetized with isoflurane (0.5%-1%) to minimize movement. A 60.96cmx34.29cm monitor was placed at a 45-degree angle and 22cm from the mouse’s head to cover 72 degrees in azimuth and 72 degrees in elevation. Green light (560 nm) was first used to focus on and image the cortical surface. Functional imaging was then performed under red light (630 nm) at 400 μm below the surface. An electron multiplying CCD camera (Cascade 512B; Roper Scientific) imaging at 30 Hz was used to capture the change in reflectance of the red light. For presenting the visual stimuli, we used a horizontal contrast-reversing checkerboard bar that was 30 degrees wide, drifting upward or downward repeating 20 times at 12 seconds/cycle. A separate recording was performed for each viewing session (binocular, contralateral eye only, ipsilateral eye only). To calculate the strength of visually driven responses, we performed a Fourier Transform of the time-series data for each pixel at the stimulus frequency (12 Hz) and computed the amplitude of the Fourier Transform using custom written python scripts (https://github.com/Palpatineli/oi_analyzer). The binocular zone was defined as the cortical region that was driven by both the ipsilateral and contralateral eye. To align the binocular region defined from optical imaging to our injection site, we matched the surface vasculature imaged under green light during optical imaging to the vasculature recorded above our imaged neuron during two-photon imaging. We then overlayed the cortical activity heatmap acquired from optical imaging.

### Analysis of Two-Photon Imaging Data

#### Preprocessing

To correct for the motion present during two-photon imaging, we concatenated the recordings for each FOV during binocular, ipsilateral eye, and contralateral eye viewing for a single timepoint. We then used suite2p (Pachitariu et al., 2016) to perform motion correction of the concatenated recordings and select regions of interests (ROIs) in the FOV. The average fluorescence intensity within the ROI was calculated for each frame, and the time-series raw fluorescence values were imported to MATLAB for subsequent analysis. An annulus around each ROI was used to estimate neuropil signal, with the neuropil corrected signal computed as *F_raw_* − 0.7 ∗ *F_neuropil_*.

#### Registration Across Multiple Timepoints

We registered neurons imaged across multiple timepoints using a MATLAB based script that aligns the average intensity projection of two timepoints using fixed coordinates (https://github.com/ransona/ROIMatchPub) with manual curation.

#### Analyzing Visual Response Properties

To normalize the neuropil corrected fluorescent values, we computed a Z score for each frame: 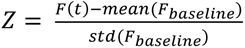, where F_baseline_ was found by concatenating all the interstimulus periods from 1 second before to the onset of each stimulus (El-Boustani et al., 2018; Jenks and Shepherd, 2020). For characterizing stimulus aligned calcium activity, we removed trials where the peak baseline activity within 1 second prior to the stimulus was above 3 standard deviations from the mean. To identify visually responsive somas, we applied three criteria for each unique stimuli: 1) at least 5/10 trials were not removed, 2) the mean amplitude response R(θ) was above 0.5 Z (which falls above the 98 percentile of responses during the interstimulus period), where R(θ) was found by subtracting the mean pre- and post-stimulus activity trace and taking the average across the 10 trials, and 3) a Student’s paired-t test between the mean pre-and post-stimulus activity trace was significant (p < 0.05) (Jenks and Shepherd, 2020). To determine the false discovery rate of our response criteria, we randomly selected timestamps from the recording as fictitious stimulus onset times and ran this permutation analysis 10,000 times. The average false discovery rate for neurons pooled across viewing sessions and days was 2.6%.

### Calculation of visual Properties

We computed a neuron’s tuning properties using a vector based approach as previously described (Mazurek et al., 2014). We calculated the orientation selectivity by the length of *L_ori_*, where 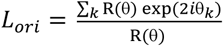. R(θ*k*) is the response at orientation *k* (taken by averaging the response across the 2 opposing directions). To determine the preferred orientation of the grating, we took the arctangent of the imaginary and real component of vector *L*, to find the corresponding angle. For assessing significant changes in the orientation preference and orientation selectivity index (OSI) of chronically tracked neurons, we performed a bootstrapped resampling of trials (10,000 times) at each of the 8 directions to generate a distribution. We then used a permutation test to determine if the mean orientation preference or OSI fell outside of the 95% percentile from the first and second timepoint.

We measured the ocular dominance index (ODI) in neurons that were visually responsive to both the contralateral eye and ipsilateral eye viewing session. We used the trial-averaged response amplitude at the neuron’s preferred direction and spatial frequency for each separate session to calculate the ODI, where *ODI* = (*C* − *I*)/(*C* + *I*). C and I refer to the contralateral eye and ipsilateral eye viewing responses.

We measured the binocular response modulation (BRM) in neurons that were visually responsive across monocular and binocular viewing sessions. We used trial-averaged response amplitude at the neurons preferred direction and spatial frequency for each separate session to calculate the BRM, where *BRM* = *B*/(*C* + *I*). For B+C and B+I cells, only the responsive monocular session’s response was used as the denominator. B, C, and I refer to the binocular, contralateral eye, and ipsilateral eye viewing responses, respectively.

To determine the alignment among the tuning preferences during the binocular, contralateral eye and ipsilateral eye viewing session, we calculated the Pearson’s correlation coefficient between the trial-averaged response amplitudes across the 8 directions at the preferred spatial frequency for each session. Orientation mismatch between viewing sessions was calculated as the absolute value of the difference in their preferred orientations.

### Linear regression model analysis

To quantify neuronal responses to binocular stimulus using responses to ipsilateral and contralateral stimulus, we first identified the preferred stimulus direction for each neuron and then selected the spatial frequency that showed the largest standard deviation of responses across stimulus directions under the binocular session. Subsequent analyses were performed using this selected frequency. The resulting directional tuning curve was fit with a von Mises function:

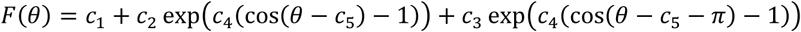

where *c*_1_ denotes the baseline response, *c*_2_ and *c*_3_ denote the amplitudes of the preferred and opposite-direction components, respectively, *c*_4_ controls tuning width, and *c*_5_ denotes the preferred direction. Model parameters were estimated by nonlinear least-squares fitting using MATLAB *lsqcurvefit* function. Responses were shifted by the absolute value of their minimum when necessary to ensure non-negative values before fitting. The fitted tuning curve was evaluated at 360 equally spaced angular positions.

We next quantified the contributions of contralateral and ipsilateral responses and their interaction to the binocular response. For each neuron, the fitted responses were normalized to [0 1] across all sessions and stimulus directions. The binocular response was then predicted as:

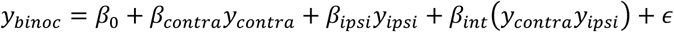

Where *y_binoc_*, *y_contra_*, and *y_ipsi_* denote the tuning curves of binocular, contralateral, and ipsilateral responses, respectively, *β*_0_ denotes the intercept, and *β_contra_*, *β_ipsi_*, and *β_int_* denote the weights for contralateral responses, ipsilateral responses, and their interaction, respectively, and *ε* denote error term.

The model was fit using a Lasso linear model (MATLAB function *lassoglm*) with an identity link and (*λ*=0.01). The regularization parameter was fixed a priori rather than optimized separately for each neuron. To estimate model performance and reduce sensitivity to a particular partition of the data, the analysis was repeated 50 times. In each repetition, data were randomly partitioned into three folds. The model was trained on two folds and evaluated on the held-out fold, with this procedure repeated across all three folds. Predictive performance was quantified as the squared Pearson correlation coefficient (*R*^2^) between the observed and predicted responses in the held-out data.

The contribution of each predictor was quantified by an ablation analysis. For each held-out dataset, one predictor at a time was set to zero while keeping the fitted model parameters unchanged. The contribution of each predictor was defined as the decrease in predictive (*R*^2^) relative to the full model:

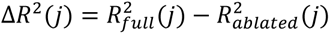

where *j* denotes the contralateral, ipsilateral, or interaction predictor. Negative values of Δ*R*^2^(*j*) were set to zero. The resulting Δ*R*^2^(*j*) values were averaged across folds and the 50 repetitions for each neuron. The regression coefficients for the contralateral, ipsilateral, and interaction terms were similarly averaged across cross-validation folds and repetitions.

### Visual Decoding Analysis

We used support vector machine (SVM) decoding with a linear kernel implemented using the libsvm library (Chang and Lin, 2011) to decode the orientation at a given trial using the trial-to-trial response amplitudes of randomly sampled neurons. For the training and testing dataset, the number of trials in each session was matched to prevent bias for training classifiers. We used 5-fold cross-validation by leaving a 20% subset of trials for prediction to avoid overfitting. This procedure was repeated 100 times. Hyperparameters such as *C* regularization weight was determined by optimization to minimize loss of validation dataset in a grid search manner (searched range 10^−3^ − 10^3^). For computing the accuracy of the SVM, we compared the decoding accuracy using empirical data with shuffled responses to normalize our decoding accuracy to chance level.

To estimate the extent to which stimulus orientation representations generalize across viewing sessions, we performed cross-session population decoding analyses. Classifiers were trained to decode stimulus orientation from neuronal activity in one viewing session (e.g., binocular) and were subsequently tested either on the same session or on the other session (e.g., ipsilateral). This procedure was repeated 100 times. To quantify changes in generalization across developmental stages (D1, D5, and D10), we defined a normalized generalization index. For each stage, decoding accuracy obtained from cross-session generalization (*ACC_ggen_*, ❷) and within-session decoding (*ACC_rrit_*_ℎ*in*_, Figure.5A, bottom, ❶) were combined to compute a normalized difference defined as (*ACC_ggen_* − *ACC_rrit_*_ℎ*in*_)/(*ACC_ggen_* + *ACC_rrit_*_ℎ*in*_) (**Figure 5A**). This metric captures the relative strength of generalization while accounting for overall decoding performance.

### Statistical Analysis

The statistical tests are detailed in the figure legends. For most analyses, we used a two-tailed non-parametric statistical analysis (i.e. Wilcoxon’s signed-rank test, Wilcoxon’s rank sum test, or Kruskal-Wallis test) or a bootstrapped permutation test repeated 10,000 times. To determine the contribution of developmental time points on response properties such as ODI and BRM or the relationship between ODI and BRM, we used linear mixed-effects models. In some cases where we had to test multiple conditions, we used a 2-way ANOVA or a one-way ANOVA and corrected for multiple comparison’s using Tukey’s post hoc HSD test or LSD test. All statistical tests were performed in MATLAB.

## Notes

### Competing Interest Statement

The authors have declared no competing interest.

